# Conformational Ensemble of RNA Oligonucleotides from Reweighted Molecular Simulations

**DOI:** 10.1101/230268

**Authors:** Sandro Bottaro, Giovanni Bussi, Scott D. Kennedy, Douglas H. Turner, Kresten Lindorff-Larsen

## Abstract

We determine the conformational ensemble of four RNA tetranucleotides by using available nuclear magnetic spectroscopy data in conjunction with extensive atomistic molecular dynamics simulations. This combination is achieved by applying a reweighting scheme based on the maximum entropy principle. We provide a quantitative estimate for the population of different conformational states by considering different NMR parameters, including distances derived from nuclear Overhauser effect intensities and scalar coupling constants. We show the usefulness of the method as a general tool for studying the conformational dynamics of flexible biomolecules as well as for detecting inaccuracies in molecular dynamics force fields.

## 1. Introduction

Many biomolecules are highly dynamic systems that undergo significant conformational rearrangements during their function. Experimental techniques such as nuclear magnetic resonance (NMR) spectroscopy, fluorescence spectroscopy and small-angle X-ray scattering (SAXS) are well-suited to probe the dynamics of molecules in solution. However, obtaining a full description of structure and dynamics of biomolecules using experiments alone can be highly non-trivial, because the measured quantities are generally time and ensemble averages over conformationally heterogeneous states. In this perspective, maximum entropy (1–3) (MaxEnt) and Bayesian (4) approaches have emerged as powerful theoretical tools for integrating simulations with experiments. Such approaches typically generate a structural ensemble for the system of interest using Molecular Dynamics (MD) or Monte Carlo simulations. This ensemble, however, may not necessarily agree with available experimental data, due to limited sampling or to inaccuracies in the employed model describing the physics and chemistry of the system (i.e. the force field). The underlying idea behind MaxEnt is to minimally perturb a simulation ensemble so as to match the experimental data. Random as well as systematic errors can be taken explicitly into account. The modification to the ensemble can be either performed on-the-fly, or even a posteriori by reweighting existing simulations. These approaches have been successfully employed to study protein systems (5), while applications to nucleic acids have been so far limited (6, 7).

In this paper we consider the conformational ensembles of four RNA tetranucleotides by integrating available NMR data (8–10) with extensive atomistic MD simulations. Despite their apparent simplicity, tetranucleotides are particularly challenging systems both from the experimental and computational point of view. First, they display significant dynamics: therefore one single structure cannot be representative of the entire ensemble. The conformational heterogeneity makes it non-trivial to provide a structural interpretation of average measurements using standard three-dimensional structure determination tools. Second, current state-of-the-art molecular dynamics force fields fail in predicting the properties of these tetranucleotides (11). Several studies (10, 12) have shown MD simulations to over-stabilize so-called intercalated conformations (see Fig.1), that in some cases correspond to the predicted free-energy minimum. From the experimental point of view the presence and the population of intercalated conformations is expected to be low, but cannot be accurately quantified.

**Fig. 1.**
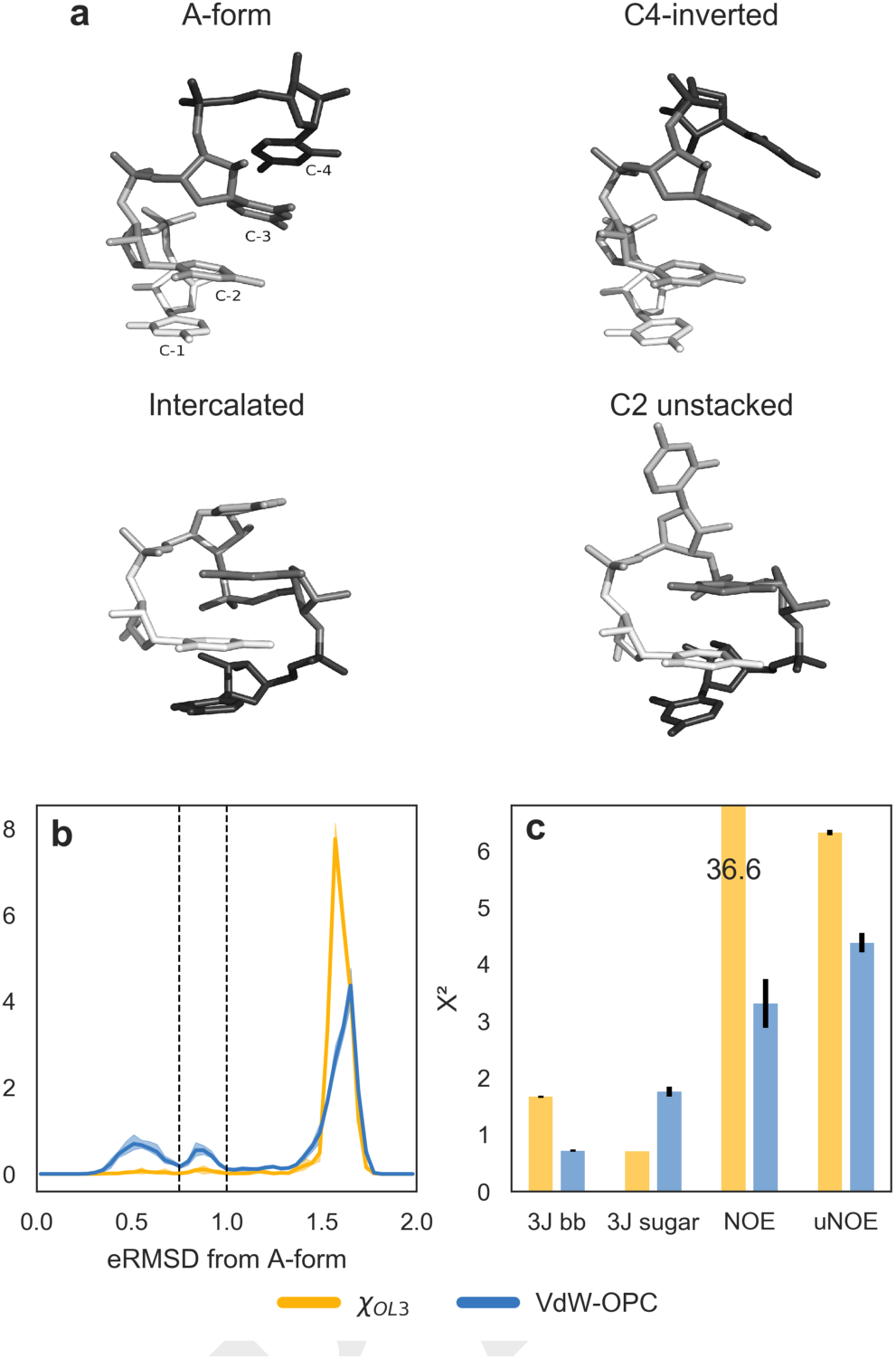
**a**) three dimensional structures of r(CCCC) discussed in the main text. **b**) eRMSD from A-form histogram for _χOL3_and _χOL3_-VdW-OPC simulations. Solid lines indicate the average calculated using a blocking procedure, while the area between minimum and maximum is shown in shade. The histogram displays three peaks corresponding to different conformations: A-form-like (eRMSD<0.75), C4-inverted (0.75 <eRMSD <1) and intercalated/C2 unstacked (eRMSD> 1.0). Thresholds are shown as dashed lines. **c**) Agreement between simulations and experiments quantified using the _χ^2^_ statistic for backbone scalar couplings (3J bb), sugar scalar couplings, NOE and unobserved NOE (uNOE). Error bars in black show the standard error of the mean.

Here we show that, even with the aforementioned complications, it is possible to obtain an accurate thermodynamic description for a system of interest by combining experiments and simulations. We report extensive atomistic MD simulations in explicit water for r(AAAA), r(CCCC), r(GACC) and r(UUUU) tetranucleotides. Except for the sequence, no other prior structural knowledge of the systems is used in simulations. We show substantial disagreement between predicted and experimental NMR data, even when using recent force-field parameters. We therefore employ the MaxEnt/Bayesian approach to refine the simulated ensembles so as to match a set of available NMR experimental data, including NOE intensities and scalar couplings. Analysis of the optimal ensembles shows that r(CCCC) and r(GACC) are mostly – but not exclusively – in A-form-like conformations. r(AAAA) and r(UUUU) display a higher complexity, as the optimal ensembles consist of a mixture of A-form with other conformationally heterogeneous structures.

## 2. Results

### A. Agreement between experiments and simulations

We first consider the tetranucleotide with sequence CCCC. NOE measurements for r(CCCC) were found to be consistent with a conformational ensemble mostly composed of A-form like structures, with a minor population (13%) of conformations with cytosine at position 4 (C4) inverted (9) (see Fig. 1a). Extensive MD simulations with the standard AMBER force field (_χOL3_ described in the Methods section) showed the presence of highly populated intercalated structures in which C1 is interposed between C3 and C4 (10, 12), while C2 is either stacked on C3 or solvent exposed. The lack of A-form-like structures is confirmed in our _χOL3_ simulations, as shown in the eRMSD histogram from ideal A-form in Fig. 1b, yellow line. To measure distances between three dimensional structures we here use the eRMSD, an RNA specific metric distance based on the relative orientation and position of nucleobases (13). It has recently been reported (14) that corrections to oxygen van der Waals radii (15) in conjunction with the OPC water model (16) (here called _χOL3_-VdW-OPC) significantly disfavor the presence of intercalated structures in r(GACC) and r(CCCC) tetranucleotides, thereby stabilizing A-form-like conformations. When using the _χOL3_-VdW-OPC force field (Fig. 1b, blue line), we observe a small, yet significant population of A-form like structures (eRMSD<0.75) as well as C4-inverted conformations (0.75-1.0 eRMSD from A-form).

The higher accuracy of _χOL3_-VdW-OPC with respect to _χOL3_ is further confirmed by the improved agreement between calculated and experimental data. Fig. 1c reports the χ^2^ for backbone ^3^J scalar couplings (H3-P, H5’/H5”-P, H4-H5’/H5”), sugar ^3^J couplings (H1’-H2’, H2’-H3’, H3’-H4’) and NOE intensities (9, 10). Additionally, we consider the absence of specific peaks in the NOESY spectra as a source of information. On the basis of assigned chemical shifts, NMR spectra were inspected for the presence of NOE cross-peaks between every pair of non-exchangeable protons in the tetramers. To assign unobserved NOEs (uNOE), the maximum NMR observable distance was estimated for each potential NOE from the minimum detectable cross-peak volume (see Methods). Whenever simulations predict a shorter distance between such proton-pairs, it is considered a violation of a uNOE. Note that the importance of unobserved NOE have been discussed for protein systems as well (17). Unobserved NOEs are of particular importance because several violations are present in intercalated structures (10). It can be clearly seen in Fig. 1c that the _χOL3_-VdW-OPC force field provides a better agreement with experimental data, especially for NOEs. We note, however, the higher χ^2^ for ^3^J sugar scalar couplings with respect to the standard _χOL3_ force field.

### B. Reweighting procedure

It is evident from Fig.1c that the conformational ensemble predicted by simulations alone is not in complete agreement with experiments. We therefore generate a conformational ensemble that satisfies the experimental constraints using the MaxEnt/Bayesian approach with the inclusion of error treatment (4, 6). In MaxEnt approaches one seeks the minimal perturbation of the simulated ensemble (i.e. the prior distribution) that satisfies a set of known experimental averages. This can be achieved (2, 6) by minimizing the function

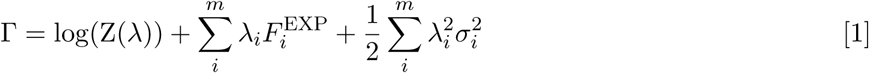

with respect to the set of Lagrange multipliers λ = λ_1_…λ_*m*_. Here, the index *i* runs over the *m* experimental averages 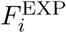 with associated normally distributed and uncorrelated errors σ_*i*_. Z is the partition function 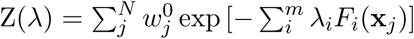 where *F_i_*(x_*j*_) is the function used to back-calculate the experimental observable from the atomic coordinates x, and 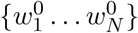 correspond to the weights of the *N* frames in the prior distribution. Note that this approach is completely equivalent to a Bayesian ensemble refinement approach (4, 18) in which one seeks the optimal weights {*w*_1_ … *w_N_*} minimizing the log posterior *L*

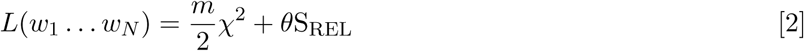

where 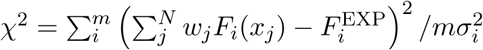 is the deviation from the experimental averages, and the relative entropy 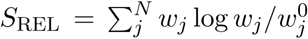 quantifies the deviation from the prior distribution. *θ* is a parameter that sets the relative weight between these two quantities, and needs to be chosen e.g. via L-curve selection.

A few items are worth highlighting. First, the number of experimental constraints, *m*, is typically much smaller compared to the number of samples, *N*, and it is therefore in practice easier to minimize the function in Eq.1 rather than Eq.2. Second, *θ* enters the MaxEnt formulation (Eq.1) as a global scaling factor of all Gaussian errors σ_*i*_. Third, heterogeneous data (NOE, ^3^J couplings, chemical shifts, etc.) can be used simultaneously in the reweighting procedure, both as averages as well as inequality constraints (6).

### C. Choosing the data and the confidence parameter

Before proceeding to the analysis of the optimized ensemble, we study the dependence of the results on i) the type of experimental data used for reweighting and ii) the tunable parameter, *θ*. Given the better initial agreement with experimental data, we here consider the _χOL3_-VdW-OPC simulations. Figure 2a shows _χ^2^_ as a function of *θ* when using scalar couplings as the only input for reweighting. As expected, small *θ* corresponds to a better fit, while in the limit of large *θ* we approach the original, unreweighted χ^2^ value (dashed line). We can also monitor the behavior of χ^2^ relative to data that were not used in the reweighting (Fig. 2a). In the limit of *θ*→ 0 the violations of uNOE become very small. Conversely, the agreement with NOE distances has a clear minimum around *θ* = 3. When using only NOEs for reweighting (Fig. 2b), we observe improved agreement with respect to all other experimental sources of data. This effect is more pronounced when using uNOE only (Fig. 2c), demonstrating the importance and the validity of this type of data. Note that, at least for r(CCCC), the reweighted χ^2^ values are always smaller compared to the original, unreweighted values, indicating that the different types of data are consistent. Given the cooperative effect of the different types of data, we finally consider the case in which ^3^J couplings, NOE and uNOE are all used at the same time for reweighting (Fig. 2d). This combination provides the best accord both for r(CCCC) as well as for the other tetranucleotides (Figs. S1-S3).

**Fig. 2.**
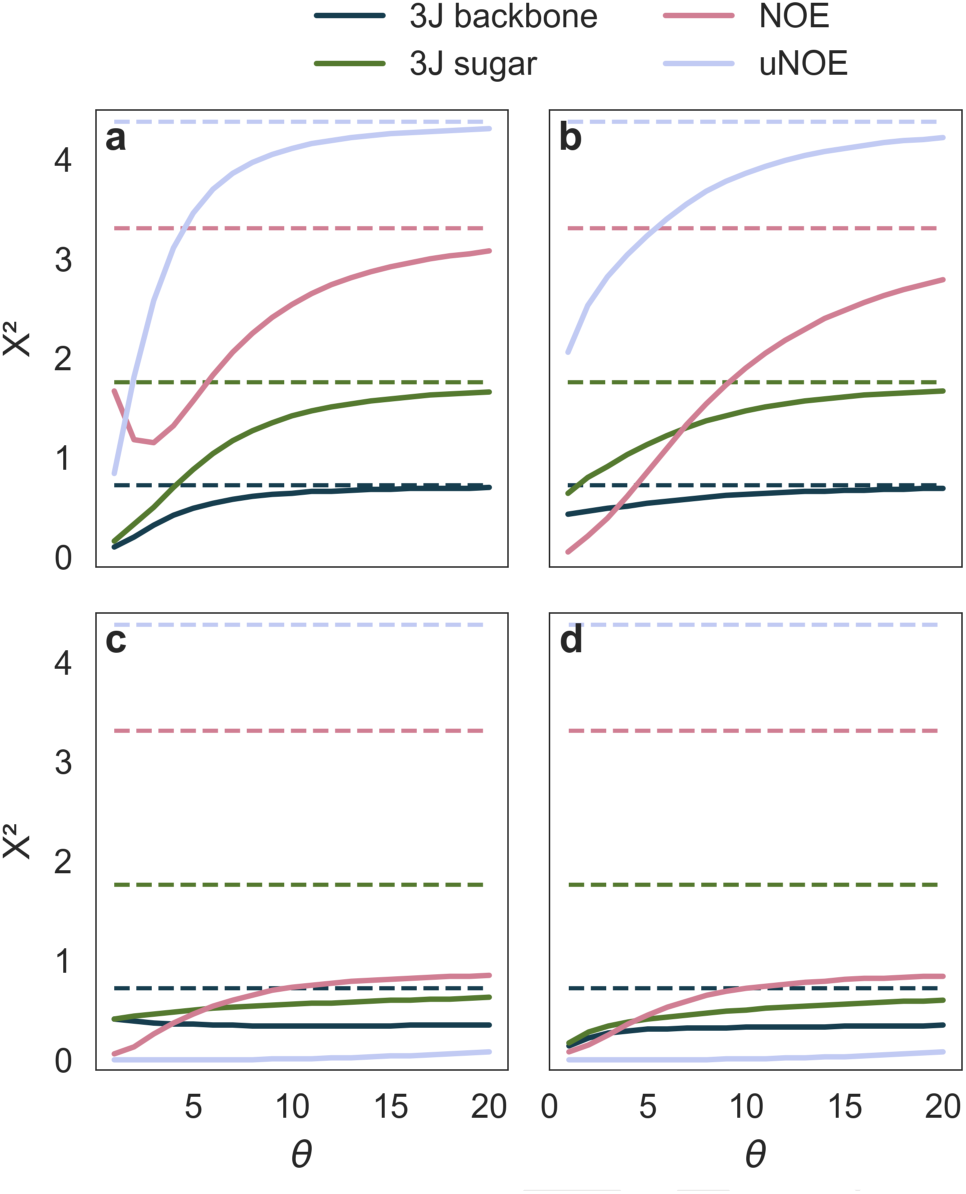
Agreement between reweighted r(CCCC) simulations and experiments as a function of the parameter *θ.* χ^2^ for different values of *θ* are reported when using **a**) scalar couplings only, **b**) NOE distances and **c**) uNOE distances. Results using all three types of data are shown in panel **d**. Initial, unreweighted χ^2^ are shown as dashed lines.

When considering χ^2^ alone one would choose a small *θ*, so as to attain the best fit. In the limit *θ* → 0, however, the original ensemble can be substantially distorted, to the point that the physico-chemical information contained in the force field is lost (Eq. 2). Additionally, this has a detrimental effect on the statistical errors, as the number of effective frames contributing to the ensemble becomes very small (Fig. S4). In order to strike a good balance between fit and proximity to the prior distribution, we scan different values of *θ* until a further decrease of this parameter leads to an increase in the relative entropy without substantially improving the fit (4). While this procedure does not provide a unique *θ*, makes it possible to identify a range of reasonable values (Fig. S4). We here use a pragmatic approach and set *θ* = 2, the largest value for which χ^2^ < 2 for all tetranucleotides and all types of experimental data. Scatter plots comparing individual experimental averages against simulations before/after reweighting are shown in Fig. S5-S8.

### D. Conformational ensemble of r(CCCC)

The set of optimized weights can be now used to calculate the full probability distribution of any observable (e.g. distances, torsion angles, etc.). In order to appreciate the properties of the optimized ensemble it is again interesting to consider the distribution of the distance from A-form (Fig. 3a). The original _χOL3_-VdW-OPC MD ensemble consists of ≈18% A-form structures (eRMSD from A-form < 0.75), and 9% of structures with C4 either inverted or unstacked (eRMSD from A-form in the 0.75-1.0 range). From the histogram of eRMSD relative to intercalated structure (Fig.3b), the initial ensemble estimates a 53% population of intercalated structures, that can be subdivided into fully stacked intercalation (13%, eRMSD < 0.4) and intercalated structures with C2 unstacked (≈ 40%, eRMSD in the 0.4-0.8 range).

**Fig. 3.**
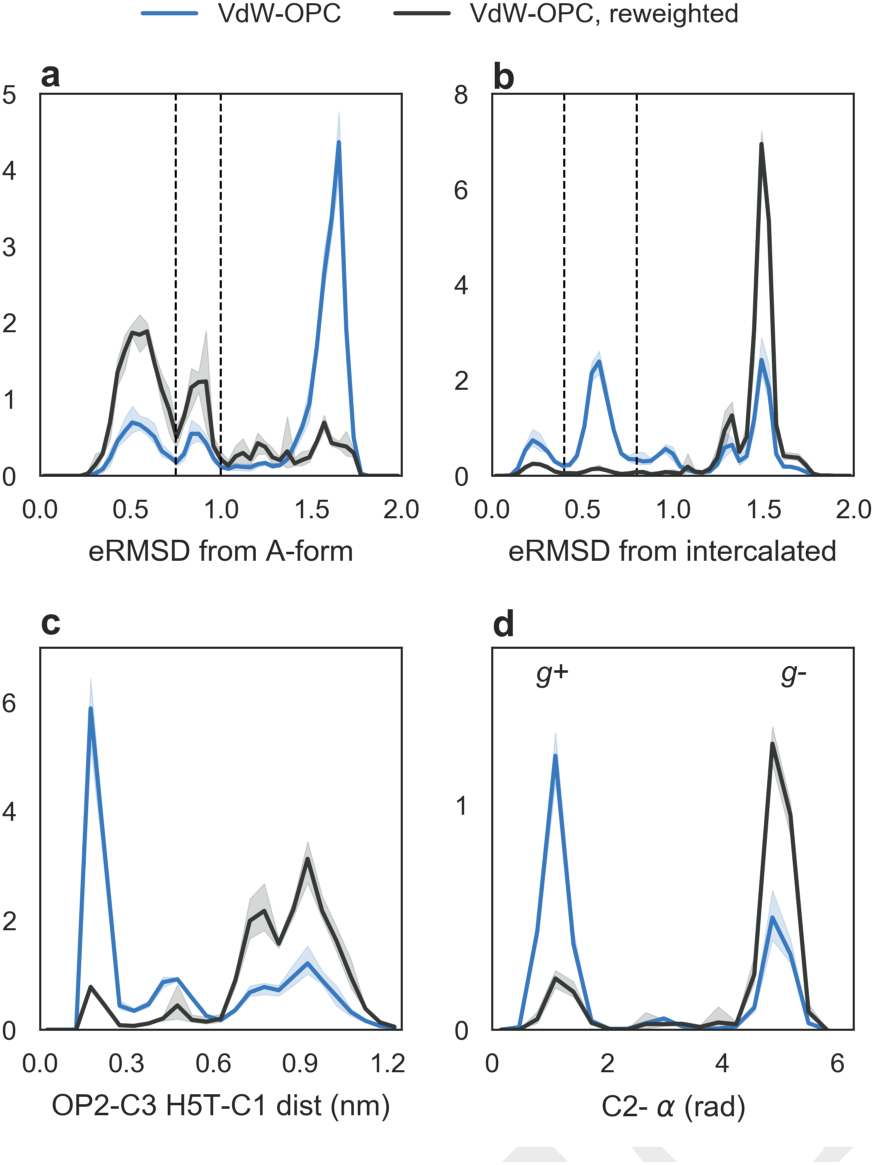
Distribution of different observables before and after reweighting r(CCCC) simulations using _χOL3_-VdW-OPC. Solid lines indicate the average calculated using a blocking procedure, minima and maxima are shown in shade. **a**) eRMSD from ideal A-form, **b**) eRMSD from an intercalated conformation, **c**) distance distribution between OP2 in C3 and H5T in C1, and **d**) the α torsion angle of C2. Peaks in panels a-b can be associated to the structures shown in Fig. 1: A-form (eRMSD from A-form below 0.75), C4-inverted (eRMSD from A-form 0.75-1.0), intercalated (eRMSD from intercalated <0.4), and intercalated with C2 unstacked (eRMSD from intercalated 0.4-0.8). eRMSD boundaries are shown as dashed lines.

Upon reweighting, A-form represents the major conformation (54%), followed by C4 inverted (22%). The population of intercalated structures is significantly reduced in the reweighted ensemble to ≈7% (Fig. 3b). This result is not surprising, as it is consistent with the picture proposed in the original experimental paper (9). The ensemble obtained here, however, did not require expert interpretation of the individual NOE distances. More importantly, the reweighting approach takes into account general properties encoded in the force-field and makes it possible to monitor degrees of freedom that were not measured by NMR. Two significant examples are reported in Fig. 3c and d. Panel c shows the distribution of the distance between the atom OP2 in C3 and the hydrogen at the 5’ terminus in C1 (H5T), where we observe the presence of a stable hydrogen bond between these two atoms (associated with the intercalated conformation) that is almost absent after reweighting. The reweighting also dramatically affects the distribution of *α* angle in C2, as we find that *gauche*^-^ (g^-^) is the preferred rotameric state in the reweighted ensemble (Fig. 3d). A similar behavior is observed for *α* in C3, *ζ* in C2 and in C3 (Fig. S10), in accordance with previous simulation studies that have shown the importance of these two torsion angles in tetranucleotides and tetraloops simulations (19, 20). We highlight that the backbone ^3^J scalar couplings used in the reweighting procedure report on *∊* and *γ* angles, but not on *α/ζ*.

### E. Conformational ensemble of r(AAAA), r(GACC), and r(UUUU)

The same procedure described above was applied to r(AAAA), r(GACC), and r(UUUU) tetranucleotides. In all cases, _χOL3_-VdW-OPC is considerably better compared to _χOL3_ force field (Fig.4, left panels). The reweighting procedure further improves agreement with experimental data. However, we do observe a residual discrepancy in some cases (χ^2^ > 1), that stems from predicted NOE distances falling outside the experimental range (Figs. S5-S8). In the case of r(GACC), three NOEs reported in the original experimental work (10) were not satisfied in a preliminary reweighting. After careful checking of the experimental data, we discovered two previously undetected spectral overlaps. The corresponding NOEs were thus removed from the list of data points. Evidently, the reweighting procedure can be used to highlight datapoints that are inconsistent with the others and, as such, might require manual inspection. These cases can be treated by using error models suitable to describe outliers (6, 21).

**Fig. 4.**
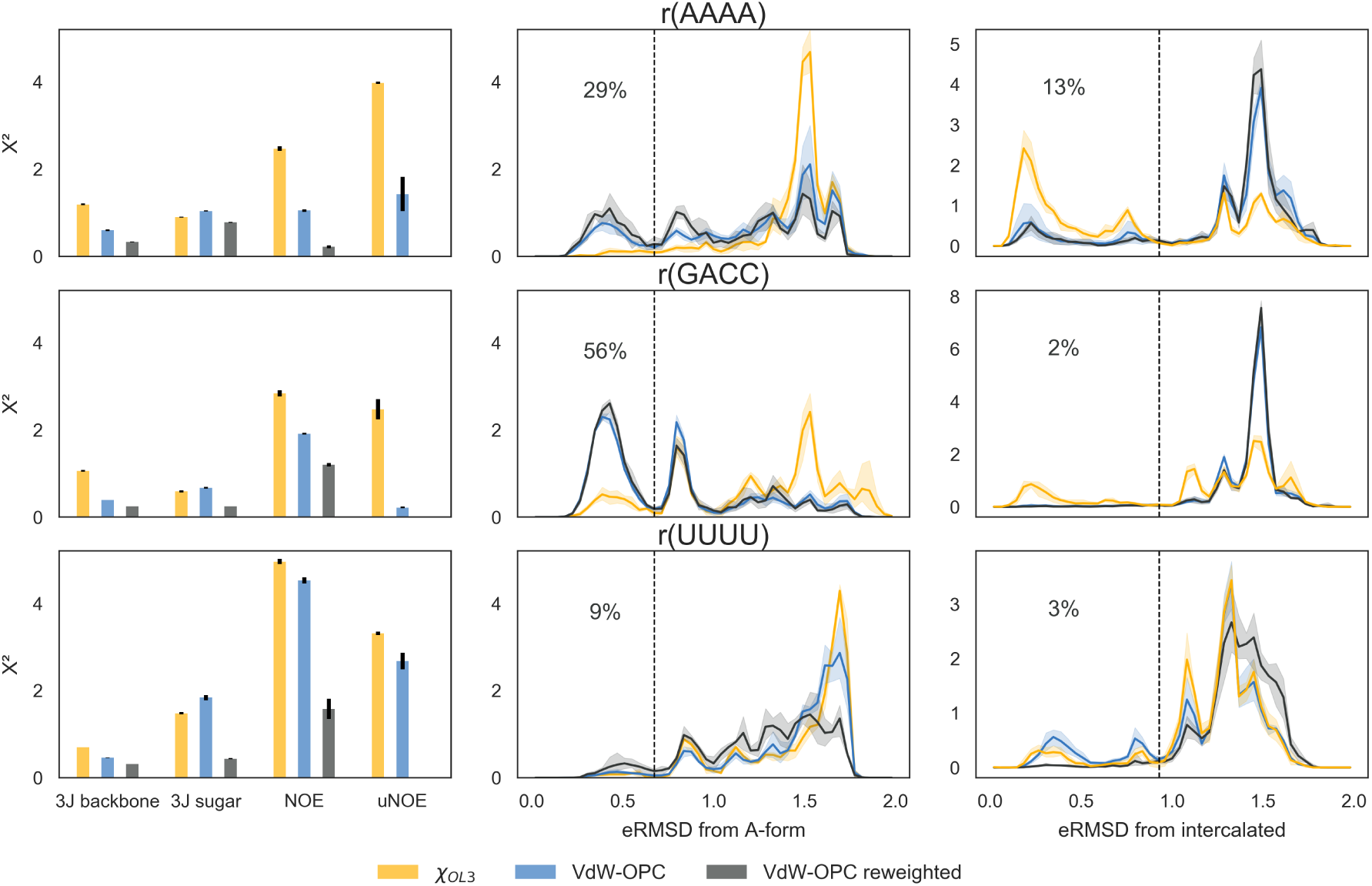
Comparison between reweighted and unreweighted ensembles for r(AAAA), r(GACC) and r(UUUU) tetranucleotides. Left panels: agreement between calculated and experimental averages for _χOL3_, _χOL3_-VdW-OPC, and reweighted _χOL3_-VdW-OPC simulations. Central panels: histogram of the eRMSD from ideal A-form. Right panels: histogram of eRMSD from intercalated. The dashed lines indicate the tresholds used for calculating the percentage of A-form-like (middle) or intercalated structures (right) upon reweighing.

The r(AAAA) ensemble is composed by ≈30% A-form like structures and 16% A4-inverted/unstacked (Fig.4, central panel). In this case, the available experimental data could not rule out completely the presence of intercalated structures, which represent the 13% of the optimized ensemble (Fig.4, right panel). The remaining 40% is composed of other structures that exhibit one or more sugar puckers in C2’-endo and/or the A1_-χ_ angle in *syn* conformation (Table 1 and Fig. S9).

**Table 1.** Percentage of C3’-endo *(δ <* 115°) and anti (χ > 120°) of reweighted _χOL3_-VdW-OPC simulations. The statistical error calculated using block averaging is below 1%.

|  | Sequence | N1 | N2 | N3 | N4 |
| --- | --- | --- | --- | --- | --- |
| % C3'-endo | AAAA | 70.6 | 75.1 | 84.5 | 66.3 |
|  | CCCC | 90.7 | 88.9 | 88.5 | 71.7 |
|  | GACC | 86.9 | 87.1 | 88.3 | 71.1 |
|  | UUUU | 61.4 | 49.5 | 50.5 | 63.2 |
| % $\chi$ Anti | AAAA | 65.2 | 96.5 | 98.4 | 97.8 |
|  | CCCC | 98.0 | 98.5 | 99.8 | 99.7 |
|  | GACC | 89.2 | 99.9 | 98.9 | 99.4 |
|  | UUUU | 88.5 | 97.1 | 96.8 | 96.3 |

r(GACC) behaves very similarly to r(CCCC), with ≈ 60%A-form-like structures and 20% of C4-inverted/unstacked. The similarity between r(GACC) and r(CCCC) can also be appreciated by considering the sugar pucker and χ angle preferences reported in Table 1 and Figs. S10-S11. Intercalation is almost completely absent in the reweighted ensembles.

Among all the systems studied here, r(UUUU) has the lowest population of A-form-like structures (9%). The rest of the ensemble is composed of a variety of diverse structures that cannot be easily clustered. This can be seen from the low percentage of sugar pucker in C3’-endo conformation (Table 1 and Fig. S12), and from the relatively flat distribution of eRMSD from A-form in Fig.4. Among this set of diverse conformations, a very small fraction of intercalated structures are present.

Note that the percentages reported here depend on two important choices: on the reference structures and on the choice of *θ.* While the geometry of the ideal A-form can be unambiguously defined (22), the intercalated structures are obtained by performing a cluster analysis of the _χOL3_ simulation as described previously (23). Although this choice has a degree of arbitrariness, we found it as a useful and intuitive manner to define an order parameter complementary to the distance from A-form. As for *θ*, we verified that the population of the different states do not depend critically on this parameter in the relevant range 2 < *θ* < 5 (Fig. S13).

## 3. Discussion

In this paper we have described the structural ensembles of four RNA tetranucleotides at the atomistic level. The characterization of these systems represents a first step in understanding the ensembles and internal dynamics of larger oligonucleotides and other RNA molecules undergoing significant conformational changes.

Due to their conformational heterogeneity, RNA oligonucleotides represent prototypical cases in which NMR experimental data need to be interpreted as ensemble averages. As such, standard procedures for NMR structure determination cannot be easily applied (24). Additionally, it is not possible to predict the properties of these systems using simulations alone, because of known force-field inaccuracies (Fig.1). Only the combination of experiment with computation makes it possible to provide an atomic-detailed description of their conformational ensembles. In this context, the MaxEnt/Bayesian approach serves as a fundamental theoretical ingredient for using the two techniques in conjunction.

In a broad sense, this can be seen as a regularization problem in which a small set of experimental data are used to gain insights into a highly dimensional, complex set of molecular conformations. The problem is under-determined, and has to be regularized by using a suitable prior distribution, here provided by MD simulations. This interpretation becomes transparent in the Bayesian ensemble refinement formulation in Eq.2 (4, 18). The balance between fit quality (χ^2^) and deviation from the prior distribution (*S*_REL_) is tuned by a system-dependent confidence parameter, *θ*, that is not known a priori. The approach used here takes explicitly into account the uncertainty *σ* on the experimental average. Since *θ* is a global scaling factor, the values of *σ* allow the relative weight of different heterogeneous data to be accounted for. Note that the calculation of the experimental observable from the atomic coordinates (i.e. the forward model) introduces inaccuracies that can be larger than the experimental uncertainty. For example, ^3^J scalar couplings calculated using Karplus relationships can introduce errors up to 2 Hz (Fig.S14). Care should be also taken when calculating NOE intensities from proton-proton distances, as the simple *r*^−6^ averaging does not take spin-diffusion into account, and it is only valid in the limit of slow internal motion compared to the tumbling time (25).

In a number of recent MaxEnt-inspired approaches a bias deriving from the experimental data is estimated on-the-fly during the simulations (4, 6, 21, 26). These approaches have the advantage of enhancing the sampling in relevant regions of the conformational space. On the other hand, the reweighting procedure can be applied a posteriori to existing simulations whenever new experimental data are available (27). Since reweighting only requires a cheap post-processing of existing trajectories, it is straightforward to perform multiple cross validation tests. Additionally, reweighting is very convenient when the forward model calculation is particularly demanding, since in biased methods the back-calculation of averages from structures has to be performed at least every few time steps (28).

In our work the reweighting approach is also used as a tool to help identify inaccuracies in molecular dynamics force fields. Modern atomistic molecular mechanics force fields consist of hundreds of parameters, and even finding the relevant interactions that can potentially improve their accuracy is a time consuming and non-trivial task. The reweighting substantially simplifies this search (Fig. 3c-d), as the probability distribution over any degree of freedom before and after reweighting can be readily compared. We find that hydrogen bonds to non-bridging oxygens are significantly destabilized upon reweighting, in accordance with previous simulation studies (10, 29). At the same time, the population of *α* and *γ* torsion angles is in some cases shifted from *gauche*^+^ to *gauche*^−^. As molecular mechanics force fields improve, the approach described here should require less experimental data to provide reliable determination of structural ensembles (30).

## Materials and Methods

### MD simulations

We have performed MD simulations on r(AAAA), r(CCCC), r(UUUU), and r(GACC) tetranucleotides. Each system was simulated with two different force-fields: i) the AMBER 99 force field (31) with parmbsc0 corrections to *α*/*γ* (32) and the _χ_OL corrections to _χ_ torsion angles (33) in TIP3P water (34). We refer to this combination as _χOL3_. These simulations were taken from our previous studies (19, 35). ii) _χOL3_with corrections to Van der Waals oxygen radii (15) and using the optimal 3-charge, 4 point (OPC) water model (16). We refer to this combination as χ_OL3_-VdW-OPC. Parameters are available at http://github.com/srnas/ff. Molecular dynamics simulations were performed using the GROMACS 4.6.7 software package (36). Ideal A-form, fully stacked initial conformations were generated using the Make-NA web server. The oligonucleotides were solvated in a truncated dodecahedric box and neutralized by adding Na+ counterions (37). Initial conformations were minimized in vacuum first, followed by a minimization in water and equilibration in NPT ensemble at 300 K and 1 bar for 1 ns. Production runs were performed in the canonical ensemble using stochastic velocity rescaling thermostat (38). All bonds were constrained with the LINCS algorithm (39), equations of motion were integrated with a time step of 2 fs. Tetranucleotides were simulated using temperature replica exchange (40) using 24 replicas in the temperature range 278 K-400 K for 1.0 *μs* per replica. All the analyses presented here were performed for the 300K replica and using 20000 frames. Averages and standard errors of the mean are calculated using four blocks of 5000 samples each.

### NMR data

Experimental NOE and scalar couplings have previously been published (9, 10). We use a Gaussian distributed experimental errors of 1.5Hz for scalar couplings (Fig. S14) and of 0.1Å for unobserved NOE. The error for NOE was estimated as min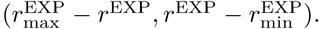The number of experimental averages for each NMR parameter and for each tetranucleotide sequence is reported in Table 2. The complete list of experimental data is available as textfiles at https://github.com/sbottaro/tetranucleotides_data. NOE intensities from simulations are calculated as averages over the *N* samples 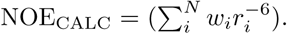 ^3^J scalar couplings are calculated using the Karplus relationships as described in Fig. S14 using the software baRNAba https://github.com/srnas/barnaba.

**Table 2.**
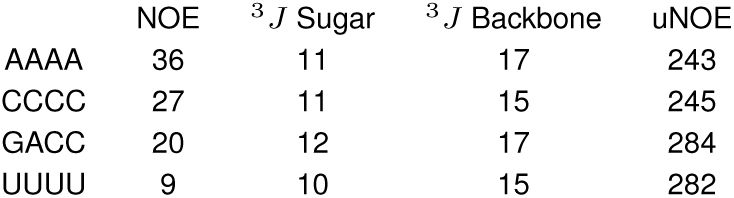
Number of experimental averages.

### Unobserved NOE

NMR spectra were inspected for the presence of NOESY cross-peaks between every pair of protons in the tetramer. If no cross-peak is observed, then the potential contact is classified as an unobserved NOE. If the spectral position of a potential cross-peak does not overlap any other observed cross-peak, then the minimum detectable cross-peak volume is assumed to be two times the standard deviation of spectral noise, *V*_err_. Scalar coupling results in NOE cross-peaks that are split into multiplets of 2, 4, or more peaks, resulting in accordingly reduced peak heights and increased minimum detectable volume. For a cross-peak consisting of *M* multiplets, the minimum detectable volume is 2*MV*_err_. *V*_err_ and a scaling factor, *c*, obtained in the original work (9, 10) from NOESY spectra with 200 msec mixing time, are used to associate a distance,R, with the minimum detectable volume: *R* = (*c*/2*MV*_err_)^1/6^. The analysis of unobserved NOEs was carried out here with 800 msec NOESY spectra where cross-peaks are typically 2.5 to 3-fold greater than at 200 msec, so the minimum detectable NOE volume was reduced by a factor of 2.5 (after correcting for any difference in number of NMR scans). If the spectral position of a potential cross-peak partially overlaps one or more observed cross-peaks, then the minimum detectable volume of the potential cross-peak is determined by the magnitude of the observed cross-peak and exact details of the overlap (instead of spectral noise). Typically, if the partially overlapped observed cross-peak is medium or weak, respectively, then a potential cross-peak exhibiting no apparent intensity is classified as unobserved with a volume that corresponds to an internuclear distance of greater than 3.3 or 4.0 A. If the overlapping observed cross-peak is strong or the potential cross-peak is close to the diagonal, then the potential cross-peak is not classified as unobserved.

## ACKNOWLEDGMENTS

The research is funded by a grant from The Velux Foundations (SB, KLL) and a Hallas-Møller Stipend from the Novo Nordisk Foundation (KLL). GB has received funding from the European Research Council under the European Union’s Seventh Framework Programme (FP/2007-2013)/ERC Grant agreement no. 306662, S-RNA-S. DHT was supported by NIH grant R01 GM22939. We thank Simon Olsson for helpful comments on the manuscript.

